# Metabolomics shows the Australian dingo has a unique plasma profile

**DOI:** 10.1101/2020.11.02.364307

**Authors:** Sonu Yadav, Russell Pickford, Robert A. Zammit, J. William O. Ballard

## Abstract

Dingoes have not been artificially selected in the past 3,500 years. They occupy a wide range of the Australian mainland and play a crucial role as an apex predator with a generalist omnivorous feeding behaviour. In contrast, humans have selected breed dogs for novel and desirable traits. First, we explore whether the distinct evolutionary histories of dingoes and domestic dogs can lead to plasma metabolomic differences. We study metabolite composition differences between dingoes (n=15) and two domestic dog breeds (Basenji n= 9 and German Shepherd Dog: GSD n=10). After accounting for within group variation, 62 significant metabolite differences were detected between dingoes and domestic dogs, with a greater number of differences in protein (n= 14) and lipid metabolites (n= 12). Most differences were observed between dingoes and domestic dogs and fewest between the domestic dog breeds. Second, we investigate variation between pure dingoes (n=10) and dingo-dog hybrids (n=10) as hybridisation is common. We detected no significant differences in metabolite levels between dingoes and dingo-dog hybrids after Bonferroni correction. However, power analyses reported that increasing the sample size to 15 could result in differences in uridine 5’-diphosphogalactose (UDPgal) levels related to galactose metabolism. We suggest this may be related to an increase in *Amylase 2B* copy number in hybrids. Our study illustrates that the dingo metabolome is significantly different from domestic dog breeds and hybridisation is likely to influence carbohydrate metabolism.

## Introduction

Natural selection leads to the accumulation of traits that are optimal for fitness and health in natural conditions as compared to artificial selection where organisms are selected for novel and desirable traits by humans. The Australian dingo and domestic dogs have experienced distinctive selection pressures. Dingoes arrived in Australia between 3,000-5,000 years ago (Savolainen et al., 2004), are ecologically, phenotypically and behaviourally distinct from domestic dogs (Smith et al., 2019), and can survive in the wild without human interference (Ballard and Wilson, 2019). The dingo maintains ecosystem balance by controlling populations of introduced mesopredators and herbivores (Letnic et al., 2012, Letnic et al., 2009). They are generalist predators and are widely distributed across mainland Australia (Doherty et al., 2019). Here, we study plasma metabolite composition differences between dingoes and two domestic dog breeds. We then investigate metabolic variation between pure dingoes and dingo-dog hybrids. In Australia, there is extensive hybridization between dingoes and domestic dog breeds (Stephens et al., 2015).

Artificial selection has led to the generation of more than 400 breeds worldwide that have a diverse range of morphological, physiological and behavioural traits (Spady and Ostrander, 2008, Wayne, 2001). We include the Basenji and the German Shepherd Dog (GSD) as representatives of domestic dogs. We selected these two breeds because the Basenji is an ancient dog breed while the GSD has an intermediate position in the current dog phylogeny and is not morphologically specialised (Parker et al., 2017) Historically, Basenjis were indigenous to central Africa and were used for hunting and guarding domestic herds (Johannes, 2004). Like dingoes, but not domestic dogs, Basenjis have an annual oestrus cycle (Fuller, 1956). GSDs are derived from common livestock dogs in continental Europe and were established as a unique breed in 1899 (Talenti et al., 2018). GSDs are a common medium to large sized domestic dog breed, bred for their intelligence and for guarding purposes (Field et al., 2020). As a result of artificial selection, specific changes have occurred in genes involved in metabolism, behaviour and development (Pendleton et al., 2018). For instance, the pancreatic amylase (*AMY2B*) copy number expansion in domestic breed dogs is considered to be an outcome of feeding on the human provided starch rich diet (Freedman et al., 2014, Arendt et al., 2016). Such dietary shifts and positive selection on metabolic genes are expected to result in differences in the metabolite profile of canids and can be quantified.

Hybridisation between dingoes and domestic dogs has occurred since European settlement in Australia (Stephens et al., 2015) and it has led to well-established morphological and coat colour variations (Smith et al., 2019). Interspecific hybrids can have an altered metabolite profile in their blood and urine likely as a result of genetic rearrangements and the difference in the metabolic pathways (Beckmann et al., 2010, Clinquart et al., 1995, Viant et al., 2009). Hybridisation is particularly common in canids with successful inter-species reproduction and survival of fertile hybrids (Gopalakrishnan et al., 2018, Gottelli et al., 1994, Galov et al., 2015, Adams et al., 2003). Such events can dilute the genetic pool of native populations and are a key threat to their genetic integrity (Gottelli et al., 1994, Roy et al., 1996). Hybridisation may not posit a threat on the genetic integrity of wild populations if its restricted to a narrow zone between geographically widespread species. However, in the case of endangered or rare species, hybridisation can lead to genetic swamping of one population by the other, disrupt adaptive gene complexes, and reduce fitness and reproductive opportunities (Rhymer and Simberloff, 1996). Here, we investigate the effects of dingo hybridisation on the plasma metabolome.

Metabolomics quantifies a large variety of small molecules from diverse pathways using biological samples and offers a direct link between organisms’ phenotypes and genotypes (Fiehn, 2002). Metabolites regulate key cellular processes such as protein activity by regulating post-translational modifications, energy source and storage, membrane stabilization as well as nutrient and cell signalling (Johnson et al., 2016). Metabolite changes are readily detectable in body fluids, and provide a more direct and meaningful biochemical interpretation as compared to other ‘omics’ techniques (van Ravenzwaay et al., 2007). An untargeted metabolomics approach detects the wide range of metabolites present in the sample without *a priori* knowledge of the metabolome composition (Johnson et al., 2016). Rapid untargeted metabolic profiling provides insights into diet associated changes in the expression of a diverse range of small molecules (Hanhineva et al., 2015). The identified metabolites (e.g., phospholipids, amino acids and vitamins) can also be used as biomarkers to inform disease progression and efficacy of clinical treatments (Khamis et al., 2017, Mamas et al., 2011, Ferlizza et al., 2020). The untargeted approach has been shown useful to discriminate inter and intra-species/breed differences in domestic dogs (Colyer et al., 2011, Lloyd et al., 2017, Beckmann et al., 2010, Carlos et al., 2020) and a single study has investigated the chemical composition in dingo scat, urine and bedding (Carthey et al., 2017). To date, no studies have explored plasma metabolite profiles in dingoes.

Blood metabolite profile between individuals and species can be shaped by genetic and by environmental factors including dietary intake, physical condition and gut microflora (Nicholson et al., 2011, Suhre and Gieger, 2012, Kettunen et al., 2012, Fujisaka et al., 2018). For instance, in several domestic dog breeds, the difference in plasma lipidome is influenced by diet under both controlled and uncontrolled dietary experiments (Lloyd et al., 2017, Boretti et al., 2020). In this study, we detected significant metabolite differences between dingoes and domestic dog breeds using a non-targeted plasma metabolome technique. Notably, dingoes differed from domestic dogs in protein and lipid metabolites. Further, metabolites related with galactose metabolism differed between pure dingoes and dingo-dog hybrids.

## Materials and methods

### Sampling and plasma preparation

To test for differences between dingoes and the domestic breeds 34 individuals were included. Ten dingoes were collected from Bargo dingo sanctuary in south-eastern Australia. Five additional dingoes from diverse geographic localities throughout Australia were included to test the generality of the results. For the domestic dogs, we included nine Basenjis from two kennels, and 10 GSDs from two kennels. All kennels were in south-eastern Australia (Table S1). The animals were between 1-10 years and closely matched for sex but unmatched on diet to keep consistency with natural conditions.

To test for differences between pure dingoes and dingo-dog hybrids a set of 10 pure and 10 hybrid dingoes collected from the same locality (Table S2). All 20 canines were aged from 1-12 years and maintained under same environmental conditions. The individuals were diet and sex matched with equal numbers of males and females. Additional samples could not be included without extreme bias of the sample design (age, purity and sex).

The purity of all dingoes and hybrid dingo status was established using the 23 microsatellite marker based dingo purity genetic test (Wilton, 2001). Basenjis and GSDs were purebred and registered with the Australian Kennel Club.

### Metabolite extraction

Blood samples were immediately stored in EDTA tubes to avoid clotting. Plasma was separated from frozen and fresh whole blood by centrifuging at 2,000g for 10 min at 4 °C. Immediately after centrifugation, plasma was transferred into clean microtubes and stored at −80 °C for further processing.

Samples were extracted following Mackay et al. (2015). Briefly, 10µl of thawed plasma samples were diluted 20-fold with cold extraction solvent (50% methanol, 30% acetonitrile, 20% water at approximately −20°C). To mix and remove any proteins, samples were vortexed for 30s, and then centrifuged at 23,000g for 10 min at 4°C. The supernatants were transferred to glass HPLC vials and kept at −80°C prior to LC-MS analysis. Pooled quality control samples were created by combining 5µL of each sample. Process blanks were created by following the extraction protocol without plasma.

LC-MS profiling was performed using Q-Exactive HF Mass Spectrometer with U3000 UHPLC system (ThermoFisher Scientific). Samples were analysed in both positive and negative heated electrospray ionization as separate injections. Samples and blanks were analysed in a random order (generated using Excel) with regular QC’s inserted into the sequence after randomisation.

A ZIC-pHILIC column (SeQuant, VWR, Lutterworth, Leics., UK) was employed to measure a broad range of metabolites of different classes as it is suggested to give the broadest coverage of metabolites with an adequate performance as compared to the other columns (Zhang et al., 2012). 5µL of the sample was injected onto the column. Separation was performed using a gradient of mobile phase A (20mM ammonium carbonate in MilliQ water, adjusted to pH 9.4 with ammonium hydroxide) and mobile phase B (100% acetonitrile) at 200µL/min. The gradient was held at 80% B for 2 minutes, ramped to 20% B at 17 minutes before returning to 80% B at 17.1 minutes and holding for re-equilibration until 25 minutes. The mass spectrometer was operated in the data dependant analysis mode – automatically acquiring MS/MS data. The instrument was scanned from 75-1000 at a resolution of 60K, with MS/MS of the top 20 ions at 15K. Source conditions were spray voltage 4.5kV positive, (3.5 kV negative), sheath gas 20 au, auxiliary gas 5 au. Heater temperature was 50°C and the capillary temperature was 275°C. S-Lens was 50V. The instrument was calibrated immediately prior to data acquisition and lock masses used to maintain optimal mass accuracy.

Data analysis was performed using Compound Discoverer software (v3.1 Thermo, Waltham, USA). The software was used to pick and integrate peaks, perform relative quantitation and attempt identification using database searches against mzCloud and Chemspider databases. The QC samples were used to correct chromatographic drift and the processed blanks used to identify and filter out background components. Before statistical analysis the data was filtered and only the most confident identifications (>50% score against mzCloud) were used. Normalised area for each metabolite was exported to excel format and then used for further statistical analysis. Metabolite classification and functions were determined using Human Metabolome Database (HMDB) and PubChem databases.

### Statistical analysis

All statistical analysis were performed in R v3.6.1 (Team and DC, 2019). An overall significant difference in the metabolites between dingoes and domestic dogs (Basenji and GSD) was determined by performing Type III ANOVA to account for within group variation.

To detect differences between the dingo, Basenji and GSD a Type II ANOVA was performed using car R package (Fox et al., 2012). Following ANOVA we obtained the pairwise difference between groups using *TukeyHSD* function in R. To identify metabolite difference between pure and hybrid dingoes a Welch two sample t-test was performed. A post-hoc power test was then performed using the *pwr*.*t*.*test* function in R (Champely et al., 2018) with a significance at P=0.05 and power of 95%. All P values obtained from statistical tests were Bonferroni (BF) corrected. All statistical analyses were performed on the combined positive ion and negative ion data sets.

## Results

### Dingo and domestic breed difference

A total of 666 metabolites were detected by LC-MS for 34 individuals. The Type III ANOVA test identified 62 significant differences between the dingo and domestic dog (Table 1). Out of 62 metabolites, a greater number of metabolite differences were detected for protein derivatives (n=14) followed by lipid derivatives (n= 12), carbohydrates (n=4) (Table 1) and others (n=32) (Table S3). Overall, the majority of proteins (71%) and lipids (66%) were lower in dingo than breed dogs while the reverse was true for carbohydrates (75%). For proteins, 11/14 metabolites were classified as amino acids and derivatives and 3/14 as peptides. The three protein metabolites that were most different between dingoes and domestic dogs (lowest P values) were Glycylglutamic acid, gamma-Glu-Gly and L-Cystine (Fig. 1A). Out of the 12 lipid differences, five were classified as phosphatidylcholines (PC) and two lysophospholipids (LyP), indicating distinction in lipid metabolism and functionality (Table 1). The three lipid metabolites with lowest P value were Linoleyl carnitine, PC (16:0/22:5n3), and Oleoylcarnitine (Fig. 1 B). The three most different carbohydrate metabolites included 1D-chiro-inositol, Istamycin C and 2,7-Anhydro-alpha-N-acetylneuraminic acid (commonly known as sialic acid) (Fig. 1C).

**Table 1:** Protein, lipid and carbohydrate differences observed between the dingo and domestic dog using type III ANOVA. Non-essential amino acid derivatives and metabolites are indicated by *.

| Broad classification | P | Subclass |
| --- | --- | --- |
| **Protein** |  |  |
| Glycylglutamic acid | 1.42E-08 | Peptide |
| gamma-Glu-Gly | 2.73E-07 | Peptide |
| L-Cystine* | 1.02E-06 | Amino acid |
| N,N-Dimethylglycine* | 1.32E-06 | Amino acid derivative |
| D-(+)-Pipicolinic acid | 2.24E-06 | Amino acid metabolite |
| 2-Amino-3-phosphonopropanoate | 3.05E-06 | Amino acid |
| Hexanoylglycine* | 4.94E-06 | Amino acid acylated |
| L-Cysteinylglycine disulfide | 7.00E-06 | Peptide |
| 4-Methylene-L-glutamate* | 1.10E-05 | Amino acid derivative |
| N-Acetyl-L-leucine | 1.25E-05 | Amino acid derivative |
| 2,6-Diaminoheptanedioic acid | 2.43E-05 | Amino acid derivative |
| N-acetyl-DL-tryptophan | 3.38E-05 | Amino acid derivative |
| N5-Ethyl-L-glutamine* | 5.76E-05 | Amino acid |
| Ophthalmic acid* | 7.38E-05 | Amino acid derivative |
| **Lipid** |  |  |
| Linoleyl carnitine | 1.51E-07 | Carnitine derivative |
| PC (16:0/22:5n3) | 2.96E-06 | Phosphatidylcholine |
| MFCD22416941/Oleoylcarnitine | 5.09E-06 | Acylcarnitine |
| PC (32:2) | 1.49E-05 | Phosphatidylcholine |
| (2E)-hexadecenoylcarnitine | 1.69E-05 | Acylcarnitine |
| PC (18:3/18:3) | 1.99E-05 | Phosphatidylcholine |
| (24R_24'R)-Fucosterol epoxide | 2.91E-05 | Epoxy steroid |
| Nervonic acid | 3.45E-05 | Fatty acid |
| LPC (22:5) | 3.93E-05 | Lysophospholipid |
| LPC 22:6 | 4.48E-05 | Lysophospholipid |
| PC (14:0/24:1) | 4.64E-05 | Phosphatidylcholine |
| PC (18:0/22:5) | 5.94E-05 | Phosphatidylcholine |
| **Carbohydrate** |  |  |
| 1D-chiro-inositol | 4.37E-06 | Sugar |
| Istamycin C | 1.62E-05 | Amino sugar |
| 2,7-Anhydro-alpha-N-acetylneuraminic acid | 4.19E-05 | Sugar |
| N-Acetylneuraminic acid | 5.51E-05 | Amino sugar |

**Figure 1:**
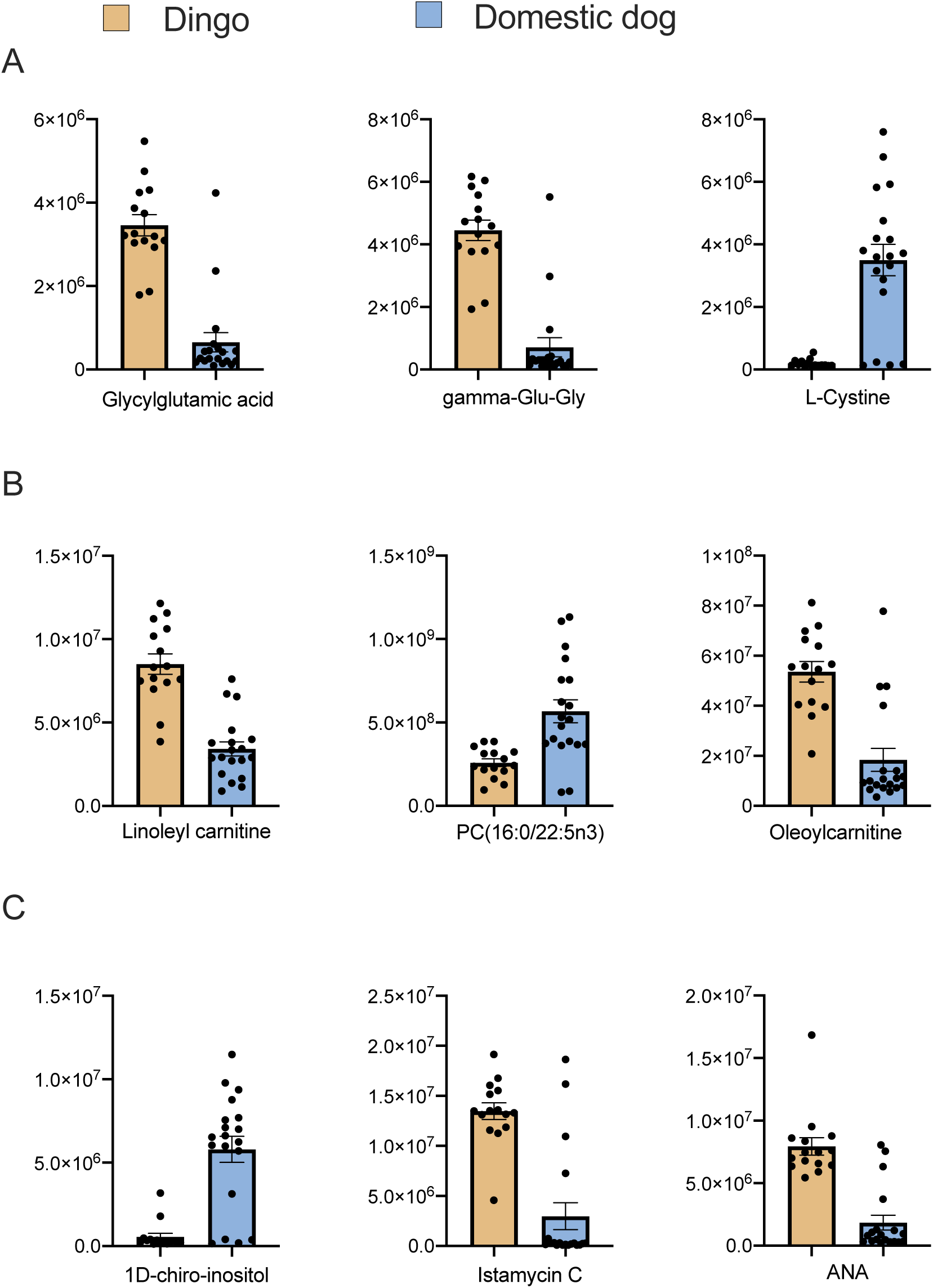
Metabolite differences between dingoes and domestic dog breeds jointly: A) Top three protein metabolite differences based on the lowest P values, B) Top three lipid metabolite differences, C) Top three carbohydrate metabolite differences. ANA: 2,7-Anhydro-alpha-N-acetylneuraminic acid (sialic acid). Y axis represents normalised area for the metabolite. Plot show mean with SE.

Overall, ANOVA showed that 98 metabolites were significantly different between dingo, Basenji, and GSD (Table S4). A greater number of metabolite differences were detected for protein derivatives (n=28) followed by lipid derivatives (n= 14), carbohydrates (n=9) and then others (n=47) (Table S4). The three most different protein metabolites were Glycylglutamic acid, gamma-Glu-Gly and N-Acetylornithine (Fig 2A). The three lipid metabolites with the greatest difference in levels were PC (18:3/18:3), 2-(2-Carboxyethyl)-4-methyl-5-pentyl-3-furoic acid, and PC (16:0/22:5n3) (Fig. 2B). The top three carbohydrate differences included Glucose-1-phosphate, UDP N-acetylglucosamine, and 1D-1-guanidino-1-deoxy-3-dehydro-scyllo-inositol (Fig 2C).

**Figure 2:**
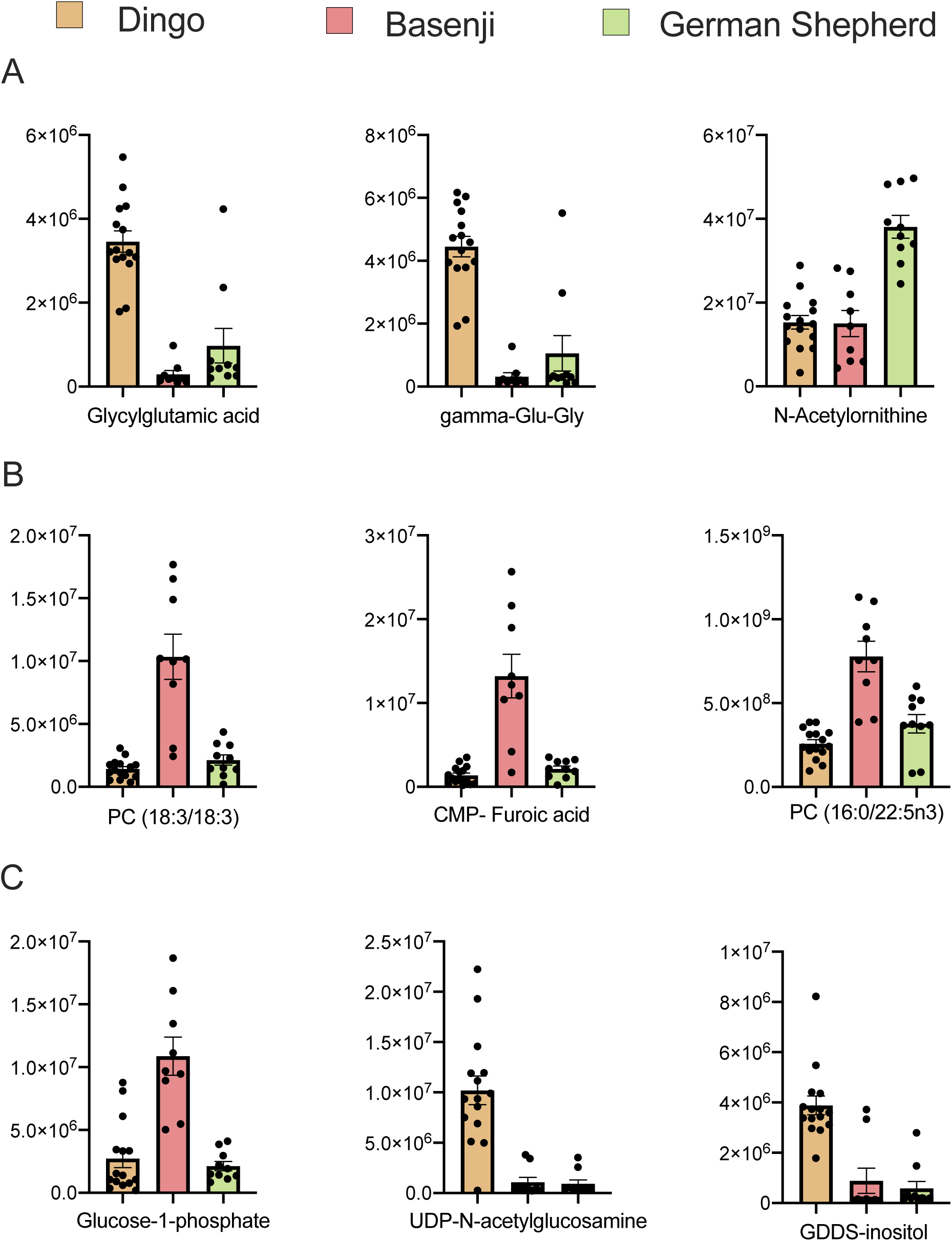
Metabolite differences between the dingo, Basenji and German Shepherd Dog: A) Top three protein metabolite differences between the three groups, B) Top three lipid metabolite differences, C) Top three carbohydrate metabolite differences. CMP-Furoic acid: 2-(2-Carboxyethyl)-4-methyl-5-pentyl-3-furoic acid, GDDS-inositol: 1D-1-guanidino-1-deoxy-3-dehydro-scyllo-inositol. Y axis represents normalised area for the metabolite. Plot show mean with SE.

Tukey’s test showed significant pairwise metabolite differences between dingoes and Basenjis (n=78), dingoes and GSDs (n=77), with fewer significant metabolite differences between Basenjis and GSDs (n=44) (Fig. 3). Between dingoes and Basenjis there were 21 unique metabolites (Table S5), 20 between dingoes and GSDs (Table S6), and no unique metabolites between Basenjis and GSDs. Comparing the dingo and Basenji, 10 lipid metabolites differed and all were lower in dingoes. In contrast, the dingo and GSD differed in 11 protein metabolites, again all lower in the dingo.

**Figure 3:**
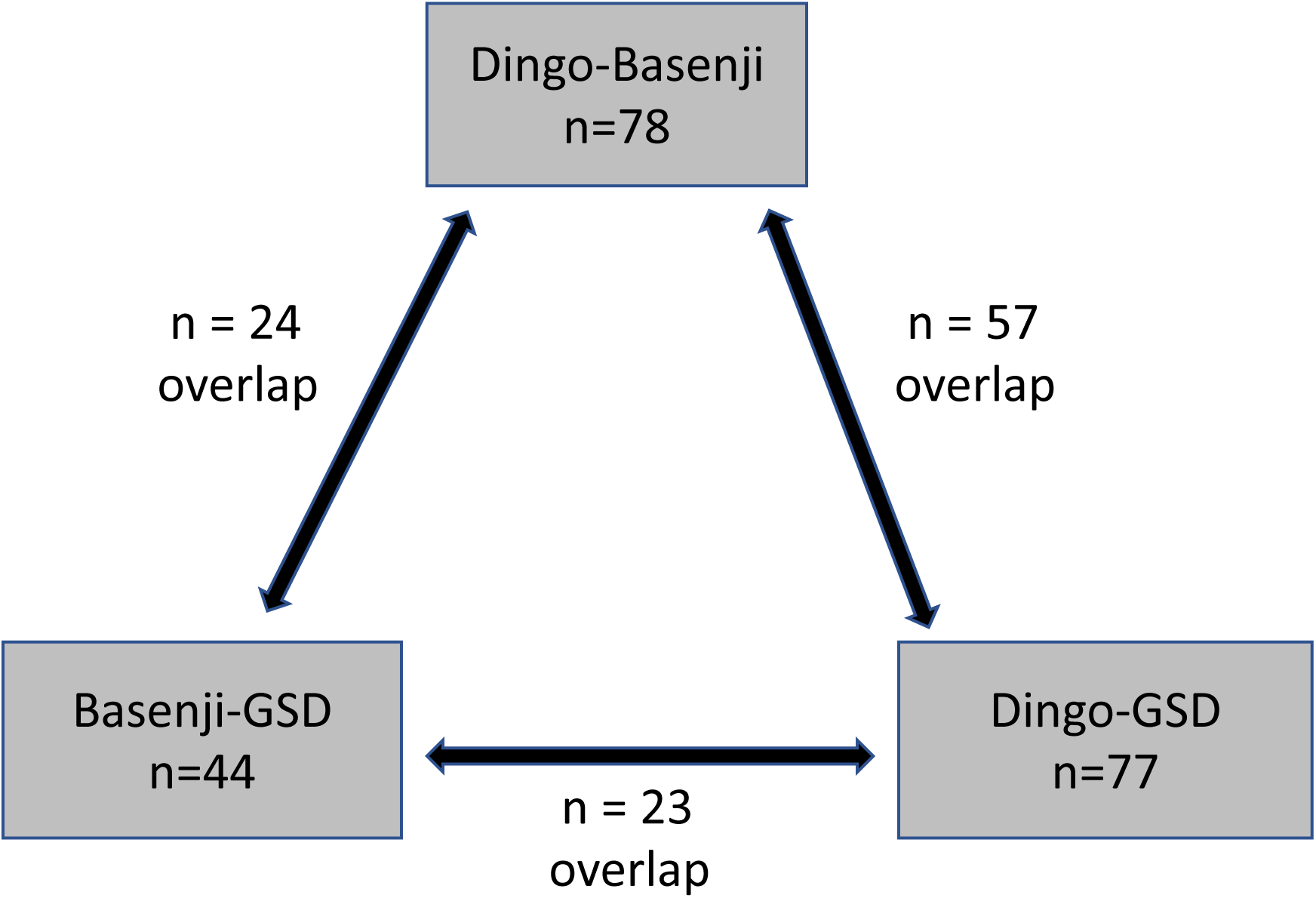
An overview of metabolite differences between the dingo, Basenji and German Shepherd Dog (GSD) detected using pairwise Tukey’s test.

### Pure and hybrid dingo differences

A total of 143 metabolites were obtained from LC-MS analysis on 10 dingoes and 10 dingo-dog hybrids. Out of these, uridine 5’-diphosphogalactose (UDPgal) (t_(17.7)_ = −3.01, P uncorrected = 0.0075), trigonelline (t_(11.03)_ = −2.37, P uncorrected= 0.037), dulcitol (t_(15.3)_ = −2.13, P uncorrected= 0.049), taurine (t_(17.52)_ = −3.73, P uncorrected= 0.002), and L-Glutathione oxidized (t_(14.46)_ = −2.33, P uncorrected= 0.03) had significantly higher levels in pure dingoes. BF correction, however, resulted in loss of significance in all cases. A post-hoc power test indicated a sample size of 15, 24 and 29 individuals respectively would result in a significant difference for UDPgal, trigonelline, and dulcitol (Fig. 4). Notably, UDPgal and dulcitol are associated with galactose metabolism.

**Figure 4:**
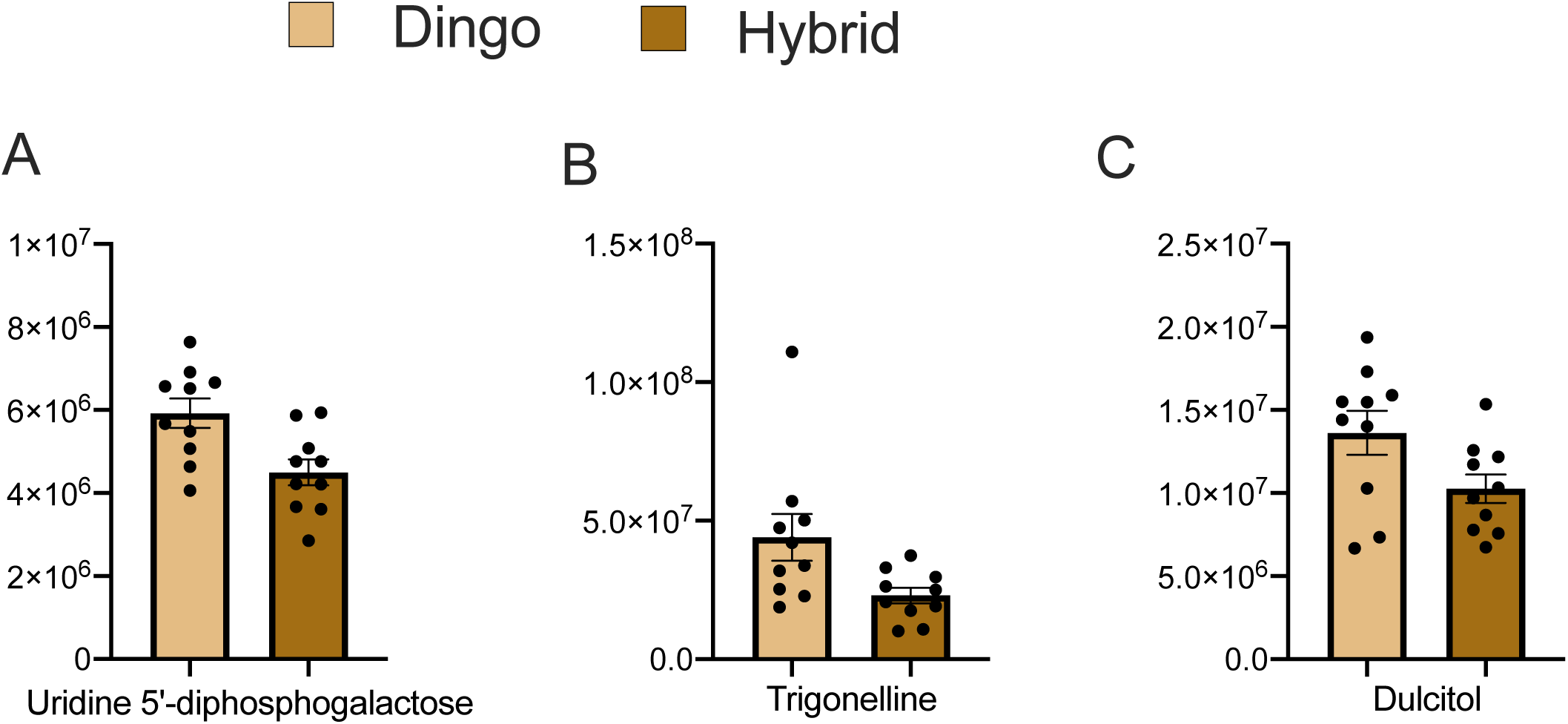
Metabolite difference between the dingo and dingo-domestic dog hybrid. Y axis represents normalised area for the metabolite. Plot show mean with SE.

## Discussion

Dingoes are Australia’s apex predator and their natural history is extensively studied (Ballard and Wilson, 2019). However, little is known about their cell biology or metabolic profile (Carthey et al., 2017). Our study reveals significant differences in the plasma metabolite composition between the dingo and domestic dogs. Of the 62 significant differences between the dingo and domestic dogs 71% of proteins and 66% of lipids were lower in dingoes. Low protein and lipid metabolite levels in dingoes may reflect genetic or dietary differences. We support the former explanation as we included dingoes and breed dogs from multiple sources. Comparing pure dingoes and dingo-dog hybrids, where animals were maintained in the same environmental conditions, metabolites associated with galactose metabolism were higher in pure dingoes. Our results provide insight into how the dingo and the domestic dog, with their distinct evolutionary histories, show variations in the cellular and metabolic pathways.

Metabolic differences involved in crucial pathways such as immune functioning and neurodevelopment indicate that the ∼8000 years of divergence of the dingo from domestic dogs have affected key genes and their metabolites essential for survival and fitness. Dingoes are generalist predators and a large proportion of the dingo diet includes protein (Doherty et al., 2019). A high protein diet may reinforce metabolites related to protein digestibility in the dingo compared to the domestic dog, which consumes food with high starch and low animal protein (Lyu et al., 2018). In our study comparing dingoes with domestic dogs, six protein derivatives that differ between dingo and domestic dogs are derived from non-essential amino acids, which are produced internally (Table 1). These protein derivative differences support our hypothesis that there are underlying genetic differences between dingoes and dogs. A recent study on the dingo reported that 50 candidate genes associated with digestion and metabolism are under positive selection (Zhang et al., 2020).

We identified multiple metabolites that are associated with neurodevelopment and likely linked with the process of domestication. The glutamate receptor agonist 2-Amino-3-phosphonopropanoate is lower in dingoes. Critically, this agonist has been shown to influence neurotransmission (Lee et al., 1995). The unsaturated fatty acid nervonic acid is also lower in dingoes than domestic dogs. Nervonic acid is tightly linked with brain development, improving memory, delaying brain aging and biosynthesis of nerve cells (Li et al., 2019). The carbohydrate sialic acid is higher in dingoes and is essential for mediating ganglioside distribution and structures in the brain (Schauer, 2000). Previously, Wang et al. (2016) showed that six genes associated with the glutathione metabolism and 49 genes associated with the neurological process and perception are under positive selection during dog domestication.

In our study, we observed significantly different levels of three protein metabolites that are associated with the bacterial community in the gastrointestinal track. Dingoes had lower levels of protein N-acetyl-DL-tryptophan, and 2,6-Diaminoheptanedioic acid and higher levels of D-pipecolic acid. N-acetyl-DL-tryptophan is a tryptophan catabolite converted by gut microbiota (Pavlova et al., 2017). It is also a protein stabilizer and protects protein molecules from oxidative degradation. 2,6-Diaminoheptanedioic acid is a lysine like derivative and is a key component of the bacterial cell wall (Webster et al., 1990). It can be found in the body fluids as a result of the enzymatic breakdown of gram-negative gut microbes. D-pipecolic acid is produced from the metabolism of intestinal bacteria (Vranova et al., 2013, Lin et al., 2018). We predict dingoes and domestic dogs will differ in their gut microbiome composition and suggest future studies explore the microbial communities in dingoes and domestic dogs raised on the same diet.

Additional metabolite differences between the dingo and domestic dog detected a suite of metabolites that influence cell signalling and immune system functioning. Of interest, the dipeptide gamma-Glu-Gly, is elevated in dingoes. Glu-Gly is an excitatory amino acid receptor antagonist in the hippocampus (Sawada and Yamamoto, 1984). L-cystine, lower in dingoes, is an oxidised form of cysteine and is linked with the immune system. L-cystine is the preferred form of cysteine for the synthesis of glutathione in immune system cells such as macrophages and astrocytes. The vitamin DL-alpha-tocopherol is lower in dingoes. It is important for regulating immune function (Lewis et al., 2019). Immune responses are expected to be higher in the dingo than the domestic dog because they are exposed to a range of environments and there is relaxed selection for high immunity in domestic dogs due to increased Veterinary intervention.

Among lipids, both LyP and all five PCs were lower in dingoes than domestic dogs (Table 1). LyP are important for cell membrane biosynthesis, energy source and storage, and intracellular signalling by acting on LPL-R lysophospholipid receptors (D’Arrigo and Servi, 2010). In addition, LyPs are involved in several fundamental processes such as reproduction, nervous system function and immunity (Birgbauer and Chun, 2006, Hla et al., 2001). PCs are the predominant component of mammalian cell membranes (Li and Vance, 2008) and are involved in the regulation of lipid, lipoproteins, and energy metabolism (van der Veen et al., 2017, Vance, 2008). Combined, the data presented in this study indicates that pure dingoes have a distinct ecological role compared to feral domestic dogs.

The study comparing pure dingoes with hybrids suggests the significant difference can be detected for UDPgal, dulcitol, and trigonelline after increasing the sample size. UDPgal and dulcitol are produced from galactose metabolism (Segal, 1995). Both metabolites are higher in pure dingoes than hybrids (Fig. 4), putatively a result of lower metabolic digestion of galactose. Potentially, this could be linked with the low *Amy2B* copy number in pure dingoes (Arendt et al., 2016). Domestic dogs are attracted to several sugars including sucrose, glucose, lactose and fructose, and have a high carbohydrate metabolic potential (Hoenig, 2014, Bradshaw, 2006). Admixture between genes from domestic dog breeds in the dingo can form new genetic combinations influencing the expression of genes involved in the carbohydrate metabolism. It is expected to result in an increasing number of *Amy2B* copies.

Future studies including East Asian breed dogs and additional hybrids will test the hypotheses presented here. Most recently, dingoes have been shown to form a monophyletic clade with East Asian breed dogs (Surbakti et al., 2020). We do not know the history of the hybrid dingoes included in this study. Including dingo-dogs hybrids with different levels of distinct domestic breeds is needed to determine whether the differences in galactose metabolism are due to increases on *Amy2B* copy number. Technically, positive controls confirming the identity of key chemical differences would strengthen our confidence in the characterization of the chemical detected.

## Conclusion

Our findings demonstrate that plasma metabolite profiling can be used to capture metabolome distinctions between the dingo and domestic dog breeds despite diet and environmental variability. Our results are consistent with the expectation that the distinct evolutionary history of dingoes and domestic dogs has played an important role in shaping pathways linked with protein, lipid and carbohydrate metabolism. A greater number of detected metabolite differences between dingoes and domestic dogs are involved in immune system functioning and neurotransmission indicating differential selection pressure on pathways crucial for fitness and survival. By comparing the pure and hybrid dingoes reared under similar environmental conditions and food, we showed that hybridisation might lead to significant differences in metabolites involved in the carbohydrate biochemical pathways.

## Acknowledgements

We thank Lucille Ellem, Ivan & Sonya Pacek, Ethel Blair, Jennifer Power, and Tony Crocker for providing samples for the study. William Donald (UNSW) provided constructive inputs. Mass spectrometric results were obtained at the Bioanalytical Mass Spectrometry Facility within the Mark Wainwright Analytical Centre of the University of New South Wales. This work was undertaken using infrastructure provided by NSW Government co-investment in the National Collaborative Research Infrastructure Scheme (NCRIS) subsidised access to this facility is gratefully acknowledged. The project was funded by Australian Research Council Discovery Project DP150102038.

## Supplementary information for

**Table S1:** Details of canines included to detect metabolite differences between dingoes and domestic breeds. NSW= New South Wales, WA= Western Australia, QLD = Queensland.

| Local ID | Group | Sex | Age | Location |
| --- | --- | --- | --- | --- |
| W0381 | Dingo | F | 4 | Bargo dingo sanctuary, NSW |
| W0380 | Dingo | M | 4 | Bargo dingo sanctuary, NSW |
| W0378 | Dingo | F | 3 | Bargo dingo sanctuary, NSW |
| W0379 | Dingo | F | 3 | Bargo dingo sanctuary, NSW |
| X3170 | Dingo | M | 8 | Bargo dingo sanctuary, NSW |
| W0235 | Dingo | M | 4 | Bargo dingo sanctuary, NSW |
| W0302 | Dingo | M | 5 | Bargo dingo sanctuary, NSW |
| W0363 | Dingo | M | 1 | Bargo dingo sanctuary, NSW |
| W0383 | Dingo | F | 3 | Bargo dingo sanctuary, NSW |
| X3172 | Dingo | M | 3 | Bargo dingo sanctuary, NSW |
| W0296 | Dingo | F | 4 | Pure dingo sanctuary, NSW |
| W0330 | Dingo | F | 4 | Pure dingo sanctuary, NSW |
| W0349 | Dingo | F | 1 | Crossroads Dingo Rescue, WA |
| W0351 | Dingo | F | >1 | RSPCA, Victoria |
| W0358 | Dingo | M | 2 | Mandurah, WA |
| BAS06 | Basenji | F | 2.5 | Basenji breed network, QLD |
| BAS07 | Basenji | M | 6.8 | Basenji breed network, QLD |
| BAS22 | Basenji | M | 6.7 | Zanzipow Basenji club, NSW |
| BAS23 | Basenji | F | 4 | Zanzipow Basenji club, NSW |
| BAS24 | Basenji | F | 4 | Zanzipow Basenji club, NSW |
| BAS25 | Basenji | F | 10 | Zanzipow Basenji club, NSW |
| BAS26 | Basenji | M | 5.7 | Zanzipow Basenji club, NSW |
| BAS27 | Basenji | F | 10 | Zanzipow Basenji club, NSW |
| BAS28 | Basenji | M | 7 | Zanzipow Basenji club, NSW |
| BAS29 | Basenji | F | 2.6 | Zanzipow Basenji club, NSW |
| GSD03 | German Shepherd | M | 2 | Allendelle Kennel, NSW |
| GSD06 | German Shepherd | F | 1.8 | Kingsvale Kennel, NSW |
| GSD07 | German Shepherd | F | 1.8 | Kingsvale Kennel, NSW |
| GSD08 | German Shepherd | F | 3.6 | Kingsvale Kennel, NSW |
| GSD 11 | German Shepherd | M | 2.2 | Kingsvale Kennel, NSW |
| GSD12 | German Shepherd | F | 2.1 | Kingsvale Kennel, NSW |
| GSD14 | German Shepherd | M | 5.6 | Kingsvale Kennel, NSW |
| GSD15 | German Shepherd | F | 4.1 | Kingsvale Kennel, NSW |
| GSD16 | German Shepherd | M | 4 | Kingsvale Kennel, NSW |
| GSD 17 | German Shepherd | F | >1 | Kingsvale Kennel, NSW |

**Table S2:**
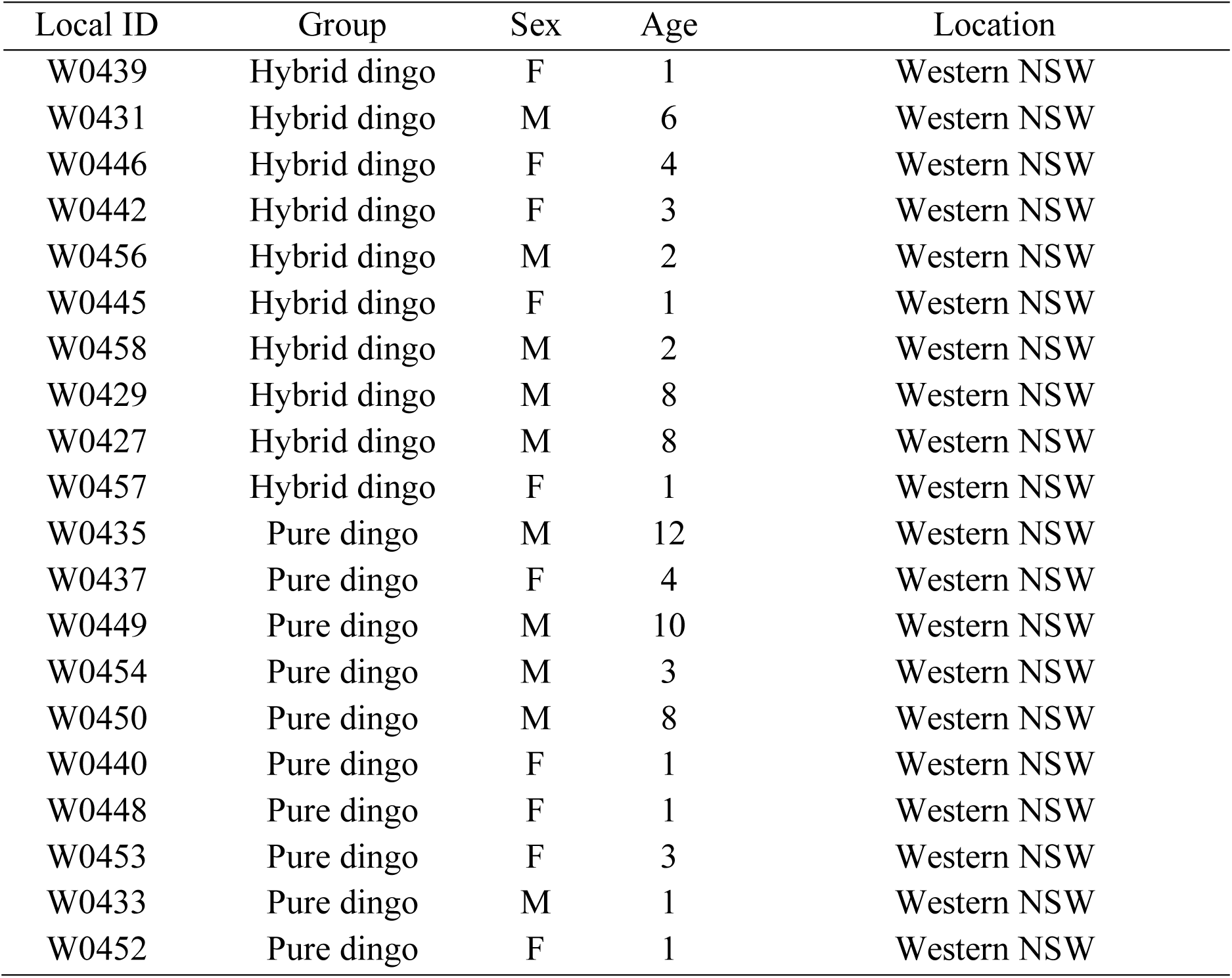
Details of dingo-dog hybrids and pure dingoes. NSW= New South Wales

**Table S3:**
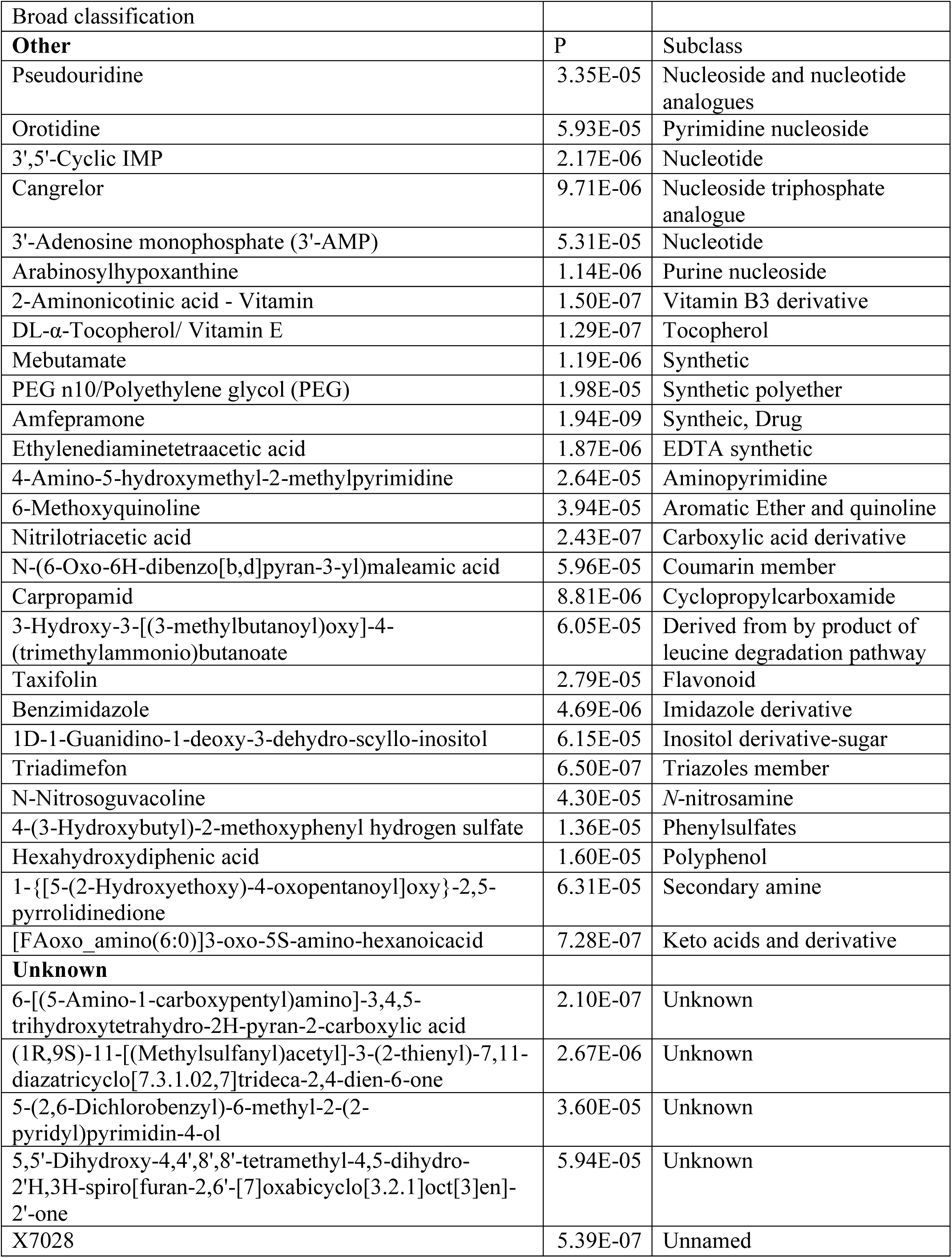
Metabolite differences between the dingo and domestic dog detected using Type III ANOVA analysis.

| Broad classification |  |  |
| --- | --- | --- |
| **Other** | P | Subclass |
| Pseudouridine | 3.35E-05 | Nucleoside and nucleotide analogues |
| Orotidine | 5.93E-05 | Pyrimidine nucleoside |
| 3',5'-Cyclic IMP | 2.17E-06 | Nucleotide |
| Cangrelor | 9.71E-06 | Nucleoside triphosphate analogue |
| 3'-Adenosine monophosphate (3'-AMP) | 5.31E-05 | Nucleotide |
| Arabinosylhypoxanthine | 1.14E-06 | Purine nucleoside |
| 2-Aminonicotinic acid - Vitamin | 1.50E-07 | Vitamin B3 derivative |
| DL-α-Tocopherol/ Vitamin E | 1.29E-07 | Tocopherol |
| Mebutamate | 1.19E-06 | Synthetic |
| PEG n10/Polyethylene glycol (PEG) | 1.98E-05 | Synthetic polyether |
| Amfepramone | 1.94E-09 | Synthetic, Drug |
| Ethylenediaminetetraacetic acid | 1.87E-06 | EDTA synthetic |
| 4-Amino-5-hydroxymethyl-2-methylpyrimidine | 2.64E-05 | Aminopyrimidine |
| 6-Methoxyquinoline | 3.94E-05 | Aromatic Ether and quinoline |
| Nitrilotriacetic acid | 2.43E-07 | Carboxylic acid derivative |
| N-(6-Oxo-6H-dibenzo[b,d]pyran-3-yl)maleamic acid | 5.96E-05 | Coumarin member |
| Carpropamid | 8.81E-06 | Cyclopropylcarboxamide |
| 3-Hydroxy-3-[(3-methylbutanoyl)oxy]-4-(trimethylammonio)butanoate | 6.05E-05 | Derived from by product of leucine degradation pathway |
| Taxifolin | 2.79E-05 | Flavonoid |
| Benzimidazole | 4.69E-06 | Imidazole derivative |
| 1D-1-Guanidino-1-deoxy-3-dehydro-scylo-inositol | 6.15E-05 | Inositol derivative-sugar |
| Triadimefon | 6.50E-07 | Triazoles member |
| N-Nitrosoguvacoline | 4.30E-05 | N-nitrosamine |
| 4-(3-Hydroxybutyl)-2-methoxyphenyl hydrogen sulfate | 1.36E-05 | Phenylsulfates |
| Hexahydroxydiphenic acid | 1.60E-05 | Polyphenol |
| 1-{[5-(2-Hydroxyethoxy)-4-oxopentanoyl]oxy}-2,5-pyrrolidinedione | 6.31E-05 | Secondary amine |
| [FAoxo_amino(6:0)]3-oxo-5S-amino-hexanoicacid | 7.28E-07 | Keto acids and derivative |
| **Unknown** |  |  |
| 6-[(5-Amino-1-carboxypentyl)amino]-3,4,5-trihydroxytetrahydro-2H-pyran-2-carboxylic acid | 2.10E-07 | Unknown |
| (1R,9S)-11-[(Methylsulfanyl)acetyl]-3-(2-thienyl)-7,11-diazatricyclo[7.3.1.0<sup>2,7</sup>]trideca-2,4-dien-6-one | 2.67E-06 | Unknown |
| 5-(2,6-Dichlorobenzyl)-6-methyl-2-(2-pyridyl)pyrimidin-4-ol | 3.60E-05 | Unknown |
| 5,5'-Dihydroxy-4,4',8',8'-tetramethyl-4,5-dihydro-2H,3H-spiro[furan-2,6'-[7]oxabicyclo[3.2.1]oct[3]en]-2'-one | 5.94E-05 | Unknown |
| X7028 | 5.39E-07 | Unnamed |

**Table S4:** Table showing 98 metabolite differences between the dingo, Basenji and German Shepherd Dog (GSD) using type II ANOVA and pairwise differences between groups obtained from Tukey’s test.

| Metabolite | P | Dingo_vs_Basenji | Dingo_vs_GSD | GSD_vs_Basenji |
| --- | --- | --- | --- | --- |
| **Protein** |  |  |  |  |
| 4-Hydroxyphenylacetyl glycine | 7.74E-06 | 8.59E-05 | 6.26E-05 | 0.998951932 |
| L.Cystine | 8.23E-06 | 0.000805194 | 1.41E-05 | 0.490665829 |
| X2.Amino.3.phosphonopropanoate | 9.38E-06 | 3.44E-05 | 0.000252339 | 0.721165446 |
| 4-Methylene-L-glutamate | 2.04E-05 | 6.81E-05 | 0.000477632 | 0.731297732 |
| L(+)-Citrulline | 3.68E-05 | 0.35783856 | 2.46E-05 | 0.004463141 |
| N5-Ethyl-L-glutamine | 4.11E-05 | 0.13272287 | 2.39E-05 | 0.017329319 |
| Hypoglycin A | 4.33E-05 | 7.58E-05 | 0.996775377 | 0.000290317 |
| Hexanoylglycine | 1.16E-05 | 0.011702382 | 8.16E-06 | 0.078400491 |
| N-Acetylornithine | 6.29E-08 | 0.996027099 | 1.81E-07 | 1.31E-06 |
| 2_6 Diaminoheptanedioic acid | 2.76E-06 | 0.222246263 | 1.73E-06 | 0.00102609 |
| L.Homocitrulline | 3.69E-06 | 0.589745147 | 3.33E-06 | 0.000296742 |
| N,N-Dimethylglycine | 4.15E-06 | 0.007140673 | 2.88E-06 | 0.057509905 |
| (3S)-3_6-Diaminohexanoate | 7.43E-06 | 0.547636654 | 6.22E-06 | 0.000610986 |
| 2.Methylserine | 1.29E-05 | 1.09E-05 | 0.006095366 | 0.089699004 |
| N-Acetyl-L-leucine | 1.83E-05 | 0.042032494 | 1.09E-05 | 0.032230509 |
| N(5)-(L-1-carboxyethyl)-L-ornithine | 3.49E-05 | 0.608533237 | 2.92E-05 | 0.001788435 |
| Ophthalmic acid | 6.35E-05 | 0.000260737 | 0.000758569 | 0.885432446 |
| D-(+)-Pipicolinic acid | 8.02E-06 | 0.000208132 | 3.15E-05 | 0.891670479 |
| 6-Acetamido-3-aminohexanoate | 2.09E-05 | 0.078925629 | 1.20E-05 | 0.018403719 |
| L-Alanyl-L-proline | 4.99E-06 | 0.891892444 | 6.75E-06 | 0.000139746 |
| Glycylglutamic acid | 8.88E-09 | 3.37E-08 | 1.90E-06 | 0.294404999 |
| gamma-Glu Gly | 9.55E-09 | 5.01E-08 | 1.19E-06 | 0.439644045 |
| L-Glutathione (reduced) | 3.69E-06 | 4.20E-05 | 3.59E-05 | 0.994421806 |
| gamma-Glu gln | 1.42E-05 | 0.622704423 | 9.86E-05 | 3.98E-05 |
| val glu | 1.98E-05 | 0.000477733 | 6.52E-05 | 0.870971035 |
| L-Cysteinylglycine disulfide | 2.61E-05 | 0.00999075 | 2.12E-05 | 0.156976629 |
| Gly Pro Glycylproline | 2.92E-05 | 0.266403734 | 1.83E-05 | 0.005607749 |
| pro gln | 5.37E-05 | 3.22E-05 | 0.201280392 | 0.00777949 |
| **Lipid** |  |  |  |  |
| Cytidine | 3.48E-06 | 4.33E-06 | 0.000963264 | 0.164723235 |
| (2E)-hexadecenoylcarnitine | 3.30E-06 | 0.000363231 | 6.38E-06 | 0.499863217 |
| MFCD22416941 | 2.35E-05 | 9.84E-05 | 0.000388888 | 0.835205992 |
| 2-(2-Carboxyethyl)-4-methyl-5-pentyl-3-furoic acid | 1.87E-07 | 2.95E-07 | 0.888601613 | 4.63E-06 |
| Diallyl adipate | 1.21E-06 | 4.81E-06 | 0.891189447 | 6.01E-06 |
| Nervonic acid | 1.69E-06 | 1.05E-06 | 0.203753158 | 0.000389254 |
| 14(Z)-Eicosenoic acid | 1.91E-06 | 2.95E-06 | 0.93026158 | 3.07E-05 |
| cis-5,8,11,14,17-Eicosapentaenoic acid | 4.99E-06 | 0.000132903 | 0.219642801 | 5.39E-06 |
| Docosahexaenoic acid | 1.05E-05 | 4.80E-05 | 0.799514938 | 3.13E-05 |
| Linoleyl carnitine | 4.63E-07 | 3.16E-06 | 1.42E-05 | 0.79647335 |
| LysoPC (22:1(13Z)) | 1.63E-06 | 3.34E-06 | 0.995669677 | 1.72E-05 |
| 1,2-di-[(9Z,12Z,15Z)-octadecatrienoyl]-sn-glycero-3-phosphocholine | 5.67E-08 | 8.36E-08 | 0.818406692 | 2.00E-06 |
| 1-hexadecanoyl-2-[(7Z,10Z,13Z,16Z,19Z)-docosapentaenoyl]-sn-glycero-3-phosphocholine | 4.00E-07 | 2.73E-07 | 0.248090621 | 8.26E-05 |
| LPC(22:6) | 2.21E-05 | 1.30E-05 | 0.167938946 | 0.004632365 |
| **Carbohydrate** |  |  |  |  |
| N-Acetylneuraminic acid | 4.00E-07 | 0.000119183 | 6.89E-07 | 0.317317543 |
| Glucose-1-phosphate | 3.18E-07 | 1.37E-06 | 0.878691883 | 1.78E-06 |
| Istamycin C | 3.09E-06 | 0.000204367 | 8.06E-06 | 0.659697612 |
| Benzoyl glucuronide (Benzoicacid) | 7.29E-06 | 0.781717133 | 8.05E-06 | 0.000284895 |
| Aminoimidazole ribotide | 3.47E-05 | 0.00141857 | 0.142093067 | 2.67E-05 |
| Uridine 5'-diphosphogalactose | 3.52E-05 | 0.000477691 | 0.000154309 | 0.970738845 |
| 1D-chiro-inositol | 1.00E-05 | 3.09E-05 | 0.000337774 | 0.643049757 |
| 2,7-Anhydro-alpha-N-acetylneuraminic acid | 2.62E-05 | 0.000423394 | 0.000108177 | 0.950466643 |
| 1D-1-Guanidino-1-deoxy-3-dehydro-scyllo-inositol | 5.14E-07 | 2.14E-05 | 2.70E-06 | 0.877081894 |
| **Other** |  |  |  |  |
| beta-Nicotinamide mononucleotide | 3.25E-06 | 7.73E-05 | 1.71E-05 | 0.942992496 |
| Pseudouridine | 6.80E-05 | 0.007362303 | 7.34E-05 | 0.35442431 |
| Cangrelor | 7.02E-05 | 0.000577774 | 0.000389073 | 0.999994364 |
| 3'-Adenosine monophosphate (3'-AMP) | 2.87E-07 | 6.15E-05 | 6.13E-07 | 0.407389452 |
| 3',5'-Cyclic IMP | 4.81E-06 | 1.36E-05 | 0.000241288 | 0.545320856 |
| Uric acid | 7.42E-05 | 4.56E-05 | 0.068308965 | 0.034132044 |
| Arabinosylhypoxanthine | 8.71E-09 | 1.45E-07 | 2.53E-07 | 0.920912441 |
| Orotidine | 8.21E-06 | 0.000480318 | 1.87E-05 | 0.648886247 |
| N4-Acetylcytidine;N-Acetyl-Cytidine | 1.58E-05 | 1.76E-05 | 0.81265826 | 0.000298371 |
| UDP N-acetylglucosamine | 4.84E-07 | 1.01E-05 | 4.54E-06 | 0.996112259 |
| DL-α-Tocopherol - Vitamin | 2.44E-06 | 2.60E-05 | 2.84E-05 | 0.981836759 |
| 2 Aminonicotinic acid - Vitamin | 1.40E-08 | 7.57E-09 | 0.001664877 | 0.000644166 |
| N-((4-AMINO-2-METHYL-5-PYRIMIDINYL)METHYL)FORMAMIDE | 4.33E-05 | 0.369376643 | 2.91E-05 | 0.004855688 |
| 4-Amino-5-hydroxymethyl-2-methylpyrimidine | 5.13E-06 | 0.218698958 | 3.17E-06 | 0.001762667 |
| 6-Methoxyquinoline | 3.38E-08 | 8.06E-08 | 0.992551967 | 5.71E-07 |
| 6-Hydroxypseudoxynicotine | 2.51E-06 | 0.657465888 | 2.49E-06 | 0.000172797 |
| D-1,2,3,4-Tetrahydroisoquinoline-3-carboxylic acid | 1.27E-05 | 0.617956701 | 1.12E-05 | 0.000760747 |
| Nitrilotriacetic acid | 2.48E-06 | 3.80E-06 | 0.000486335 | 0.224327011 |
| Carpropamid | 6.47E-06 | 1.08E-05 | 0.000726789 | 0.312753332 |
| Nitrendipine | 1.14E-06 | 1.15E-05 | 1.75E-05 | 0.953528123 |
| Amfepramone | 1.52E-11 | 1.00E-09 | 3.43E-10 | 0.992159831 |
| Dimetridazole | 5.56E-07 | 5.47E-06 | 0.528480577 | 1.39E-06 |
| Propamocarb | 2.16E-06 | 0.970446103 | 6.31E-06 | 2.17E-05 |
| Ethylenediaminetetraacetic acid | 4.13E-06 | 1.95E-05 | 0.000104162 | 0.776963989 |
| Taxifolin | 3.99E-05 | 5.89E-05 | 0.002742279 | 0.367511144 |
| 8-Hydroxyalanylclavam | 4.42E-05 | 3.21E-05 | 0.020184768 | 0.076069401 |
| 5-Nonyl-2-oxotetrahydro-3-furancarboxylic acid | 1.84E-05 | 1.15E-05 | 0.248428052 | 0.002435442 |
| Benzimidazole | 1.79E-06 | 7.64E-05 | 6.98E-06 | 0.820353547 |
| Methylimidazoleacetic.acid | 1.25E-07 | 0.792591801 | 6.90E-07 | 1.01E-06 |
| [FAoxo_amino(6:0)]3-oxo-5S-amino-hexanoicacid | 1.19E-07 | 0.003567317 | 6.76E-08 | 0.005863915 |
| 2-Amino-5-[2-(4-formylphenyl)hydrazino]-5-oxopentanoic acid | 5.91E-05 | 0.000386701 | 0.000432653 | 0.985539878 |
| N.Nitrosoguvacoline | 5.66E-07 | 7.02E-07 | 0.000324815 | 0.103479511 |
| 4-(3-Hydroxybutyl)-2-methoxyphenyl hydrogen sulfate | 2.38E-05 | 3.12E-05 | 0.002601555 | 0.278201571 |
| X3.4.Methoxyphenyl.propyl.hydrogen.sulfate | 3.46E-05 | 2.55E-05 | 0.471927573 | 0.001669411 |
| Hexahydroxydiphenic acid | 2.14E-05 | 0.000231309 | 0.000132199 | 0.998957001 |
| 1-Amino-1-deoxy-scylo-inositol 4-phosphate | 1.52E-05 | 0.001136197 | 2.63E-05 | 0.538383024 |
| 1-{[5-(2-Hydroxyethoxy)-4-oxopentanoyl]oxy}-2,5-pyrrolidinedione | 1.86E-06 | 2.99E-05 | 1.57E-05 | 0.998823833 |
| Mebutamate | 4.99E-06 | 0.000156316 | 1.91E-05 | 0.86279313 |
| N-(3,5-Dichlorophenyl)-N'-ethylthiourea | 1.07E-05 | 0.195227498 | 6.41E-06 | 0.0037323 |
| Maleic hydrazide | 4.59E-05 | 0.001618216 | 9.50E-05 | 0.711421166 |
| PEG n10 | 1.71E-05 | 0.067539046 | 9.78E-06 | 0.018632454 |
| Triadimefon | 1.27E-06 | 5.80E-06 | 4.90E-05 | 0.684905095 |
| 5,5'-Dihydroxy-4,4',8',8'-tetramethyl-4,5-dihydro-2'H,3H-spiro[furan-2,6'-[7]oxabicyclo[3.2.1]oct[3]en]-2'-one | 2.61E-08 | 4.58E-08 | 0.897229466 | 7.54E-07 |
| (1R,9S)-11-[(Methylsulfanyl)acetyl]-3-(2-thienyl)-7,11-diazatricyclo[7.3.1.02,7]trideca-2,4-dien-6-one | 2.85E-07 | 1.06E-05 | 1.86E-06 | 0.923751101 |
| 5-(2,6-Dichlorobenzyl)-6-methyl-2-(2-pyridyl)pyrimidin-4-ol | 4.60E-07 | 1.03E-05 | 4.08E-06 | 0.991315512 |
| 6-[(5-Amino-1-carboxypentyl)amino]-3,4,5-trihydroxytetrahydro-2H-pyran-2-carboxylic acid (non-preferred name) | 1.60E-06 | 5.07E-05 | 7.97E-06 | 0.904041259 |
| X7028 | 4.02E-06 | 2.18E-05 | 8.60E-05 | 0.829557536 |

**Table S5:** A total of 21 unique metabolite differences were observed between the dingo and Basenji using type II ANOVA. 4

| Broad classification | P | Subclass |
| --- | --- | --- |
| **Lipid** |  |  |
| cis-5,8,11,14,17-Eicosapentaenoic acid | 0.0001329 | Fatty acid |
| Diallyl adipate | 4.81E-06 | Fatty acid |
| Docosahexaenoic acid | 4.80E-05 | Fatty acid |
| Nervonic acid | 1.05E-06 | Fatty acid |
| 14(Z)-Eicosenoic acid | 2.95E-06 | Fatty acid |
| 2-(2-Carboxyethyl)-4-methyl-5-pentyl-3-furoic acid | 2.95E-07 | Fatty acid |
| LysoPC (22:1(13Z)) | 3.34E-06 | Lysophospholipid |
| LPC (22:6) | 1.30E-05 | Phosphatidylcholine |
| PC (18:3/18:3) | 8.36E-08 | Phosphatidylcholine |
| PC (16:0/22:5n3) | 2.73E-07 | Phosphatidylcholine |
| **Protein** |  |  |
| Hypoglycin A | 7.58E-05 | Amino acid |
| Pro-Gln | 3.22E-05 | Dipeptide |
| **Carbohydrate** |  |  |
| Aminoimidazole ribotide | 0.00141857 | Carbohydrate |
| Glucose-1-phosphate | 1.37E-06 | Carbohydrate |
| **Other** |  |  |
| Uric acid | 4.56E-05 | Purine derivative |
| N4-Acetylcytidine | 1.76E-05 | Pyrimidine nucleoside |
| Dimetridazole | 5.47E-06 | Drug |
| 3-(4-Methoxyphenyl)propyl hydrogen sulfate | 2.55E-05 | Phenylsulfates |
| 5-Nonyl-2-oxotetrahydro-3-furancarboxylic acid | 1.15E-05 | Gamma butyrolactones |
| 6-Methoxyquinoline | 8.06E-08 | Aromatic ether |
| 5,5'-Dihydroxy-4,4',8',8'-tetramethyl-4,5-dihydro-2'H,3H-spiro[furan-2,6'-[7]oxabicyclo[3.2.1]oct[3]en]-2'-one | 4.58E-08 | Unknown |

**Table S6:** A total of 20 unique metabolite differences were observed between the dingo and German Shepherd Dog using type II ANOVA.

| Broad classification | P | Subclass |
| --- | --- | --- |
| **Protein** |  |  |
| gamma-Glu-gln | 9.86E-05 | Dipeptide |
| Gly-Pro(Glycylproline) | 1.83E-05 | Dipeptide |
| L-Alanyl-L-proline | 6.75E-06 | Dipeptide |
| (3S)-3_6-Diaminohexanoate | 6.22E-06 | Amino acid derivative |
| 2_6-Diaminoheptanedioic acid | 1.73E-06 | Amino acid derivative |
| N5-(L-1-Carboxyethyl)-L-ornithine | 2.92E-05 | Amino acid |
| N5-Ethyl-L-glutamine | 2.39E-05 | Amino acid analogue |
| 6-Acetamido-3-aminohexanoate | 1.20E-05 | Beta-amino acids |
| L (+)-Citrulline | 2.46E-05 | Amino acid |
| L-Homocitrulline | 3.33E-06 | Amino acid derivative |
| N-Acetylornithine | 1.81E-07 | Amino acid |
| **Carbohydrate** |  |  |
| Benzoyl glucuronide (Benzoicacid) | 8.05E-06 | Carbohydrate |
| **Other** |  |  |
| N-(3,5-Dichlorophenyl)-N'-ethylthiourea | 6.41E-06 | Synthetic |
| PEG n10 | 9.78E-06 | Synthetic polyether |
| Propamocarb | 6.31E-06 | Drug |
| 4-Amino-5-hydroxymethyl-2-methylpyrimidine | 3.17E-06 | Pyrimidine |
| 6-Hydroxypseudooxynicotine | 2.49E-06 | Aryl alkyl ketones |
| D-1,2,3,4-Tetrahydroisoquinoline-3-carboxylic acid | 1.12E-05 | Carboxylic acid |
| Methylimidazoleacetic acid | 6.90E-07 | Imidazolyl carboxylic acids |
| N-((4-Amino-2-methyl-5-Pyrimidinyl) methyl) formamide | 2.91E-05 | Amino pyrimidine |

## Notes

### Competing Interest Statement

The authors have declared no competing interest.

